# Combining historical agricultural and climate datasets sheds new light on early 20^th^ century barley performance

**DOI:** 10.1101/2022.09.14.507767

**Authors:** Joanna Raymond, Ian Mackay, Steven Penfield, Andrew Lovett, Haidee Philpott, Stephen Dorling

## Abstract

Barley (*Hordeum vulgare ssp. vulgare*) is cultivated globally across a wide range of environments, both in highly productive agricultural systems and in subsistence agriculture and provides valuable feedstock for the animal feed and malting industries. However, as the climate changes there is an urgent need to identify adapted spring barley varieties that will consistently yield highly under increased environmental stresses. In this research we combined recently released historical weather data with published early 20^th^ century Irish spring barley trials data for two heritage varieties: *Archer* and *Goldthorpe*, following an analysis first published by Student in 1923. Using linear mixed models, we show that interannual variation in observed spring barley yields can be partially explained by recorded weather variability. We find that whilst *Archer* largely yields more highly, *Goldthorpe* is more stable under wetter growing conditions, highlighting the importance of considering growing climate in variety selection. Furthermore, this study demonstrates the benefits of access to historical trials and climatic data and the importance of incorporating climate data in modern day breeding programmes to improve climate resilience of future varieties.

## 1 Introduction

Spring barley (*Hordeum vulgare ssp. vulgare*) is the most widespread spring crop in Ireland and approximately 120,000 ha are sown each year (TEAGASC 2020). It has been grown in Ireland since the 1800s and is well suited to the Irish soils and long growing season, which offer high yield potential (TEAGASC 2017). As the climate changes and extreme weather events become more frequent, identification of spring barley varieties that prosper and consistently produce high yields is a priority.

Within barley’s germplasm there are genotypes that can tolerate abiotic stresses such as drought and heat (Bindereif *et al*. 2021; Ivandic *et al*. 2000; Wu *et al*. 2017). Barley landraces can also grow well in biogeographical zones with reduced soil fertility in which modern elite barley varieties fail to reach maturity (Schmidt *et al*. 2019). As environmental stresses become more frequent and there is a need for varieties demanding fewer resource inputs, heritage varieties may provide valuable genetic variation and a possible resource for these wild-type traits.

A well-documented set of spring barley trials data for 1901-1906 exists, comparing two heritage spring barley varieties: *Archer* and *Goldthorpe. Archer* is a 2-row narrow-eared variety that originated in East of England and outperformed the long-running favourite *Chevalier* in yield, quality and straw strength (Hunter 1913). *Goldthorpe* is a 2-row wide-eared barley known for its high malting quality. In 1889 a single wide ear was found in a field of *Chevalier* near Goldthorpe, Yorkshire and was selected and propagated to become *Goldthorpe* (Gothard *et al*. 1983; Malcolm 1983; Reid *et al*. 1929).

Analysis of these trials data by William Gosset in Student (1923) concluded that the chief difficulty in comparing variety performance was that differences between varieties are small compared with variations due to weather. Whilst weather was recorded during this period at various locations across Ireland, these data were not accessible to Student at that time.

On the approach to its centenary, Student’s 1923 paper “On testing of Cereal Varieties” remains noteworthy. It was published in the early days of establishing methods of variety testing and lists reasons why yield trials are necessary which have not changed – environmental conditions “evoke different responses in strains”; “the soil on which plants are grown is never uniform”; and “the effects of soil and weather are far greater than the differences [between varieties] which we have to investigate”.

A recent data rescue project has extended the temporal coverage of digitally available daily maximum and minimum air temperature and rainfall observations back to include this early 20^th^ century period (Mateus *et al*. 2020; Ryan *et al*. 2020). Historical climate data has been shown to be valuable for identifying the climatic influence on crops (e.g. Kahiluoto *et al*. 2019; Lopes 2022; Rezaei *et al*. 2015; Trnka *et al*. 2010) as well as the impact of the interaction between genotype and the environment (GxE) on yield (e.g. de los Campos *et al*. 2020; Fabio *et al*. 2017). Despite this, few studies take advantage of the benefits of combining historical crop trials data with historical weather data in barley (Gillberg *et al*. 2019).

In this study we show that by combining spring barley trials data with climate data, interannual variation in early 20^th^ century spring barley yields can be partially explained by recorded weather variability. We demonstrate the relative stabilities of *Archer* and *Goldthorpe* varieties and show the importance of considering the growing climate in variety selection. We also explore a range of variable selection methods and modelling tools to identify the most suitable modelling techniques for highly correlated, multi-dimensional yield and climate data. Finally, we discuss the benefits of access to trials data and the importance of incorporating climate data in modern day breeding programmes to improve the climate resilience of future varieties.

## 2 Materials and methods

### 2.1 Datasets

The barley trials dataset analysed by Student (1923) consists of two spring varieties – *Archer* and *Goldthorpe* – in unreplicated 2-acre plots at 18 distinct farm locations across the barley-growing districts in Ireland (Figure 1a and 1b). Locations for each trial site are recorded by the town and district, from which a latitude and longitude has been estimated. The number of trial sites increases each year from 4 in 1901 to 12 in 1906. Yield data was recorded in barrels and stones per acre and price was recorded in £sd per acre. To give the values modern context, these have been converted to tonnes/ha and £/ha, respectively.

**Figure 1:**
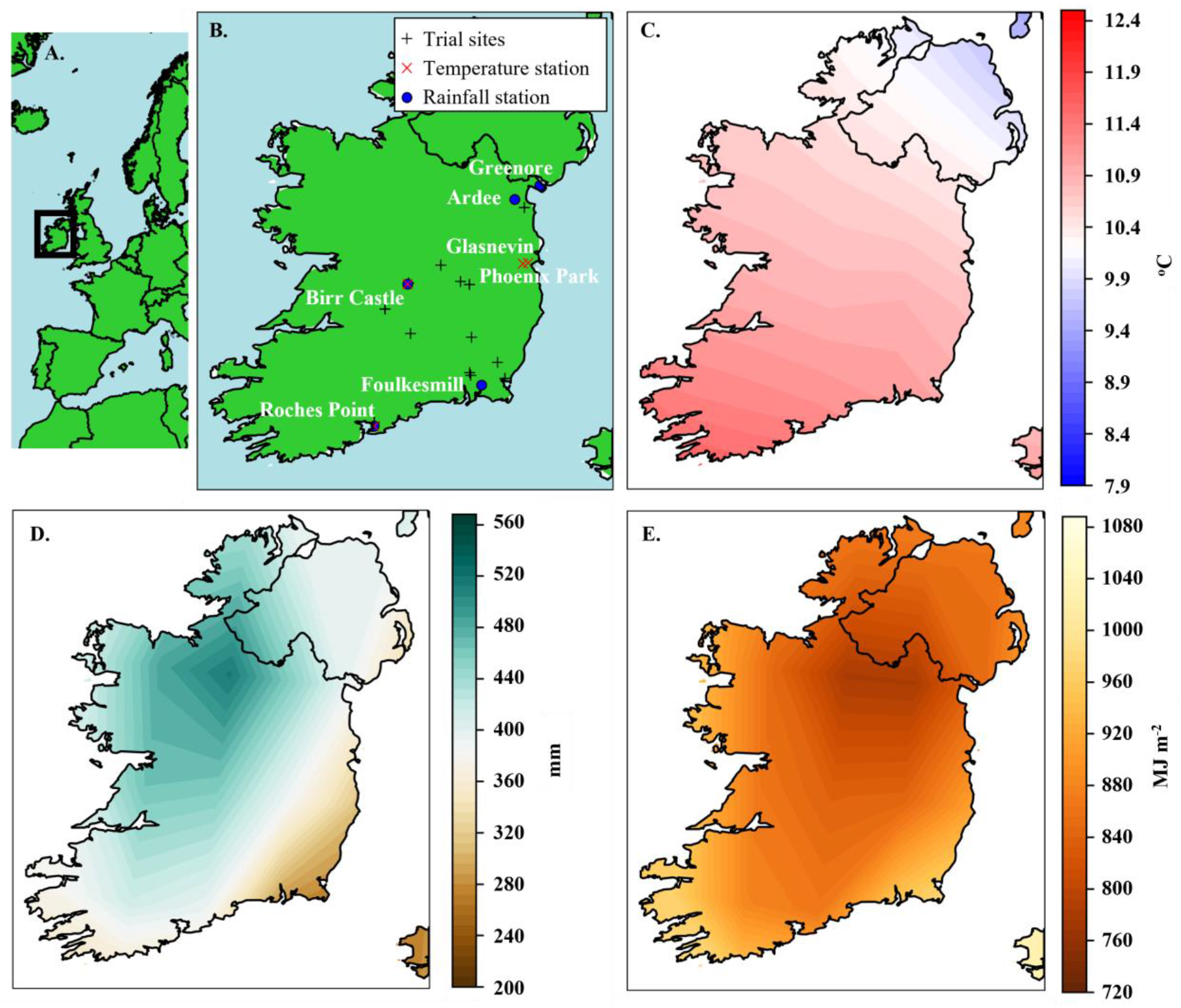
1901-1930 growing season climate for Ireland. A. The location of Ireland relative to Europe and North Africa. B. Weather stations open between 1901-1906 and closest to barley trial sites (+). Stations with rainfall only (blue circle), temperature only (red x) and both rainfall and temperature data are shown. Growing season (March-August) average temperature (°C) (C.) total rainfall (mm) (D.) and surface photosynthetically active radiation (MJ m^-2^) (E.) for 1901-1930, calculated using ERA-20C (Poli et al. 2016).

Each trial site was paired with weather data for the growing season from the nearest weather station open during the period (Figure 1b). Here growing season is defined as 1^st^ March to 31^st^ August, based on current spring barley growing practices. Daily temperature data was obtained from the Ireland Long-term Maximum and Minimum Air Temperature dataset (ILMMT) (Mateus *et al*. 2020). Daily rainfall data was sourced from Met Éireann and forms part of Ireland’s pre-1940 rainfall records (Ryan *et al*. 2020).

To enable longer-term analysis of national rainfall trends, monthly rainfall totals for the Island of Ireland (IOI) were accessed from a 305-year (1711-2016) rainfall data record (Murphy *et al*. 2018). Daily climate data for post-1960 were also sourced from the Met Éireann website (https://www.met.ie/climate/available-data/historical-data) and combined with the pre-1960 data for long-term localised climate data analysis.

In addition to station data, ECMWF’s twentieth century reanalysis dataset (ERA-20C) (Poli *et al*. 2016) was used to add a gridded and regional context to the weather stations and as a further quality control check. ERA-20C is a gridded dataset spanning 1900-2010, with a horizontal resolution of approximately 125km. Daily, invariant and monthly mean data are available: http://apps.ecmwf.int/datasets/data/era20c-daily/. Here we used the Monthly Means of Daily Means for 2 metre temperature (K) and total precipitation (m). Monthly Means of Daily Means for photosynthetically active radiation at the surface (J m^-2^) were also downloaded to supplement the lack of daily or monthly location-specific solar radiation measurement data.

Growing season average temperature (Figure 1c), total rainfall (Figure 1d) and total photosynthetically active radiation (PAR) (Figure 1e) for 1901-1930 confirm that the driest and sunniest region is the south-east of Ireland.

Growing degree days (GDD) were calculated using daily temperature data for growing season months March to August and the following equation:

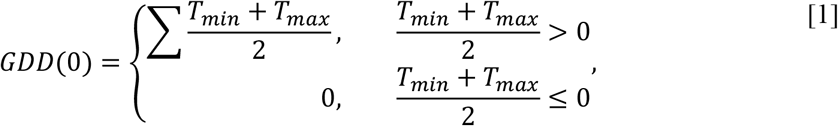

where *T_min_* is daily minimum temperature and *T_max_* is daily maximum temperature. *GDD*(0) gives a day-by-day sum of the number of degrees by which the mean temperature exceeds 0°C (Baron *et al*. 2012; Bauer *et al*. 1992; Juskiw *et al*. 2001). To ensure that missing data did not incorrectly reduce the final GDD value, growing seasons with missing data were dropped. Roches Point was missing data from 15 years (10%), Glasnevin was missing 8 years (5%) and Birr Castle no years. None of the years in 1901-1906 study period were missing data.

In the climate analysis two averaging periods are used: the full record of the dataset in question, to provide context for extreme events, and the relevant 30-year period, as the basis for the calculation of anomalies.

### 2.2 Modelling Irish spring barley and climate data

The spring barley variety trials are located across different sites, creating a clustered dataset where trial yields are not independent and not all farms were used each year. At a given site, the yields are all dependent on the same environmental factors such as rainfall and soil type, as well as the same farmer and agronomy. Therefore, the following linear mixed-effect model was used so that both farm and year could be included as random effects, using REML through *lmer* from ***lme4*** (Bates *et al*. 2020) in R:

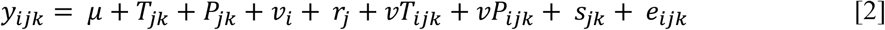

*y_ijk_* is the yield of variety *i* in year *j* at farm *k, μ* is the overall trial series mean, *T_jk_* is the effect of monthly temperature in year *j* at farm *k, P_jk_* is the effect of monthly precipitation in year *j* at farm *k*, *v_i_* is the effect of variety *i, r_j_* is the effect of year *j*, with year included as a factor variable, *vT_ijk_* is the interaction between variety *i*, monthly temperature *T_ijk_* in year *j* at farm *k*, and *vP_ijk_* is the interaction between variety *i* and monthly precipitation *P* in year *j* at farm *k* and *e_ijk_* is the residual term. *S_jk_* is the effect of site within years, representing the interaction between year term *r_j_* and farm term *f_k_*. This term means each farm is treated as different each year, which is a more accurate representation given the exact location of fields is unknown and may have varied.

The monthly variables *T_jk_* and *P_jk_* encompass the growing season (March-August) monthly precipitation and temperature. The site term *S_jk_* and year term *η* are fitted as random effects. The two genotype-by-environment terms (GxE) variety × temperature *vT_ijk_* and variety × rainfall *vP_ijk_* terms are fitted as fixed effects as the specific reaction of individual varieties (genotype) with the climate covariates (environment) is of interest.

#### 2.2.1 Variable selection methods

To reduce the dimensionality of the data and identify the most significant monthly temperature and precipitation variables in determining yield to include in [2], best subset selection, forwards and backwards stepwise selection, the lasso (Tibshirani 1996) and elastic net (Zou & Hastie 2005) were used on the linear model run using *lm* in R:

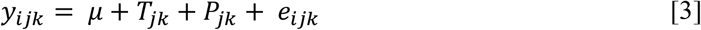

These were implemented in R using the functions and arguments detailed in Table S1. Significant variables (p < 0.05) in each of the selected models were identified using an analysis of variance (ANOVA). For each method, the RMSE and adjusted R^2^ were calculated for the selected model.

Mixed-effect model backwards elimination was also carried out using *step* on equation [2] modelled using *lmer* from ***lmerTest*** package (Kuznetsova *et al*. 2017).

#### 2.2.2 Principal Component Analysis

A Principal Component Analysis was implemented using the *ggcorr* function from ***GGally*** (Schloerke *et al*. 2021) and *prcomp* function from ***stats***.

#### 2.2.3 Pearson’s correlation analysis

Pearson’s correlation analysis was used to identify the climate covariates with the highest correlation with yield as well as the degree of correlation between the climate covariates.

#### 2.2.4 Akaike Information Criterion

Each climate variable was input into equation [2] iteratively and the significance of that variable and accompanying model Akaike Information Criterion (AIC) was calculated. The AIC was then compared with [2] without any climate variables to see if the additional variable improved the fit, using *anova(model1,model2*) in R. Additional climate variables were iteratively added and their significance and model AIC inspected.

### 2.3 Comparison of standard error of difference between means

Student (1923) calculates the standard error of the mean difference in variety means. To understand if the models run in this analysis can improve on this value, this was first repeated using the equation 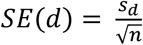, where *s_d_* is the standard deviation of the differences and *n* is the number of paired trials. After checking this against Student (1923), the value was then converted to t/ha.

To find an estimate of the standard error of difference between the varieties in the selected model, the *emmeans* function in R was used. The model and variable of interest, variety *v_i_*, were specified. The *contrast* function was then applied to this, using *method = “pairwise”*. This calculates the estimate of difference, standard error, degrees of freedom, *t*. ratio and p value for the variety pair. The statistical significance of the difference in mean values was then checked by calculating the t-statistic.

## 3 Results

### 3.1 Spring barley yields show high variation

Median yields varied from year-to-year by up to 50% for *Archer* and up to 58% for *Goldthorpe* (Figure 2). For both varieties the lowest yields occurred in 1903 (combined mean 2.2 t/ha), and highest in 1905 (combined mean 3.2 t/ha). There was large variation in yields within years, in particular for *Archer* in 1903 (SD = 0.59 t/ha) and for *Goldthorpe* in 1901 (SD = 0.67 t/ha). This was despite the number of trials increasing each year.

**Figure 2:**
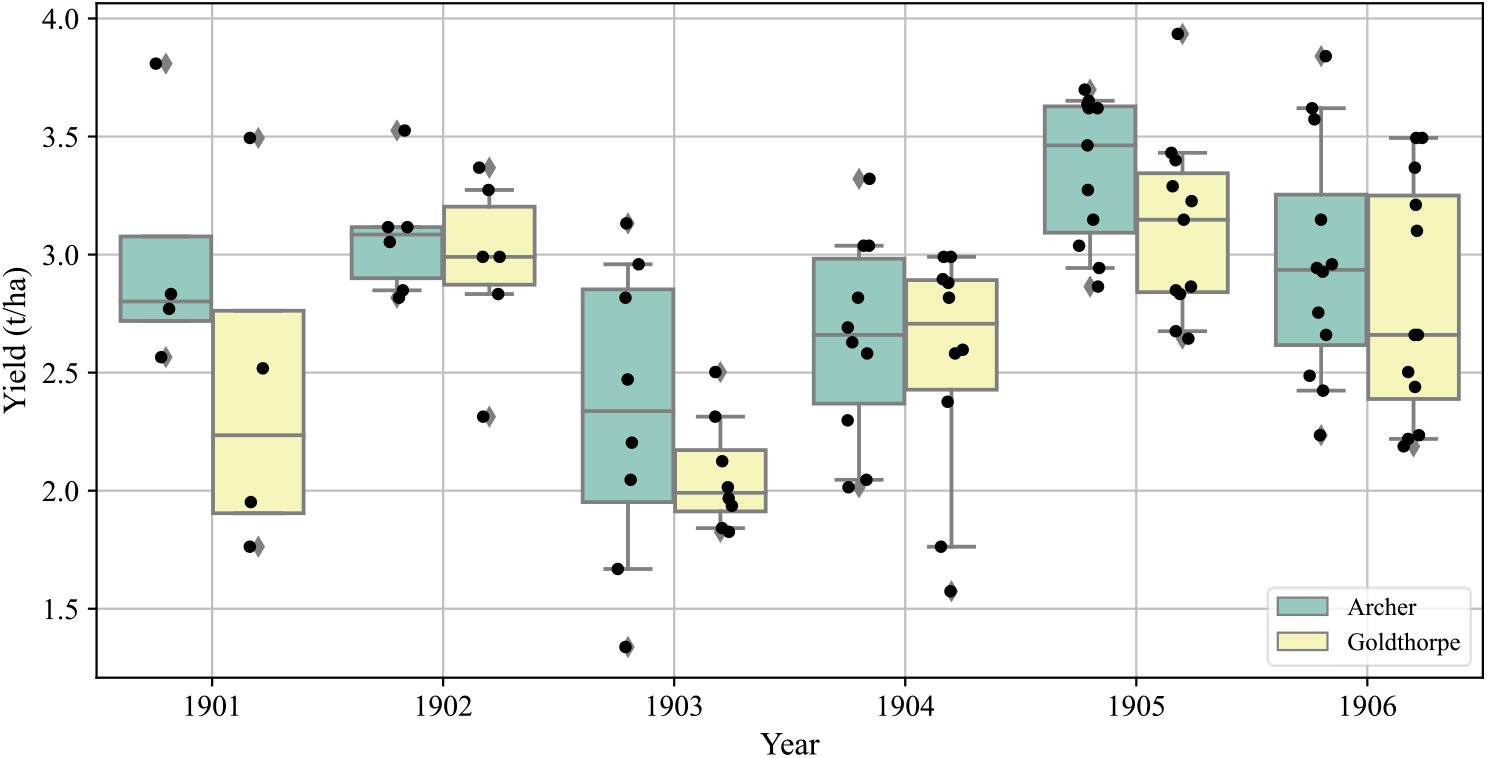
Barley trials yields. (t/ha) (black dots) for 51 trials across 18 farms between 1901-1906, for two varieties: Archer (blue) and Goldthorpe (yellow). Outliers (diamonds) represent trial yields in the 5^th^ and 95^th^ percentiles. There were 51 trials per variety, increasing from 4 in 1901 to 12 in 1906. Data from Student (1923).

Only three farmers were involved in all 6 years of the trials: Hawkins, McCarthy and Wolfe (Figure 3). There were clear differences from farm to farm in yields reflecting the differences in climate, soil type, topography, farm management practices and years. All three farms showed similar interannual variability: 1903 was the lowest yielding year while 1902 and 1905 were the highest. Yields fluctuated by up to 50% with average yields increasing ~45% (1.8 t/ha) between 1903 and 1905, indicating low stability in these varieties.

**Figure 3:**
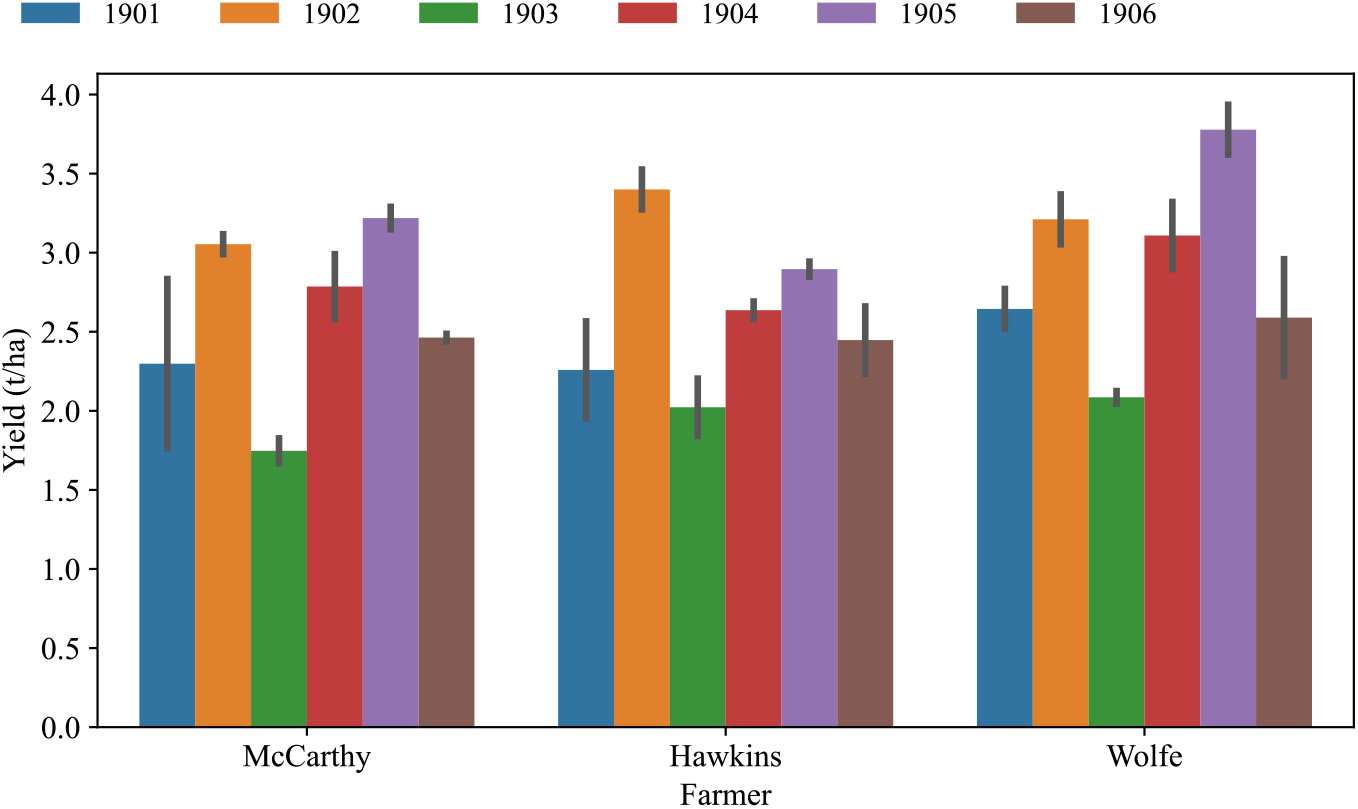
Interannual spring barley yield (t/ha) variability for three farms with data for the entire period 1901-1906. Error bars show the difference between Archer and Goldthorpe variety yields. Data from Student (1923).

### 3.2 Spring barley price shows similar variation to yield

Student used price as a measure of quality of the crop. The lowest quality of both varieties occurred in 1903 and highest in 1905 (Figure 4), as with yield (Figure 2). Student (1923) acknowledged that the value of the crop per acre was mostly dependent on the yield.

**Figure 4:**
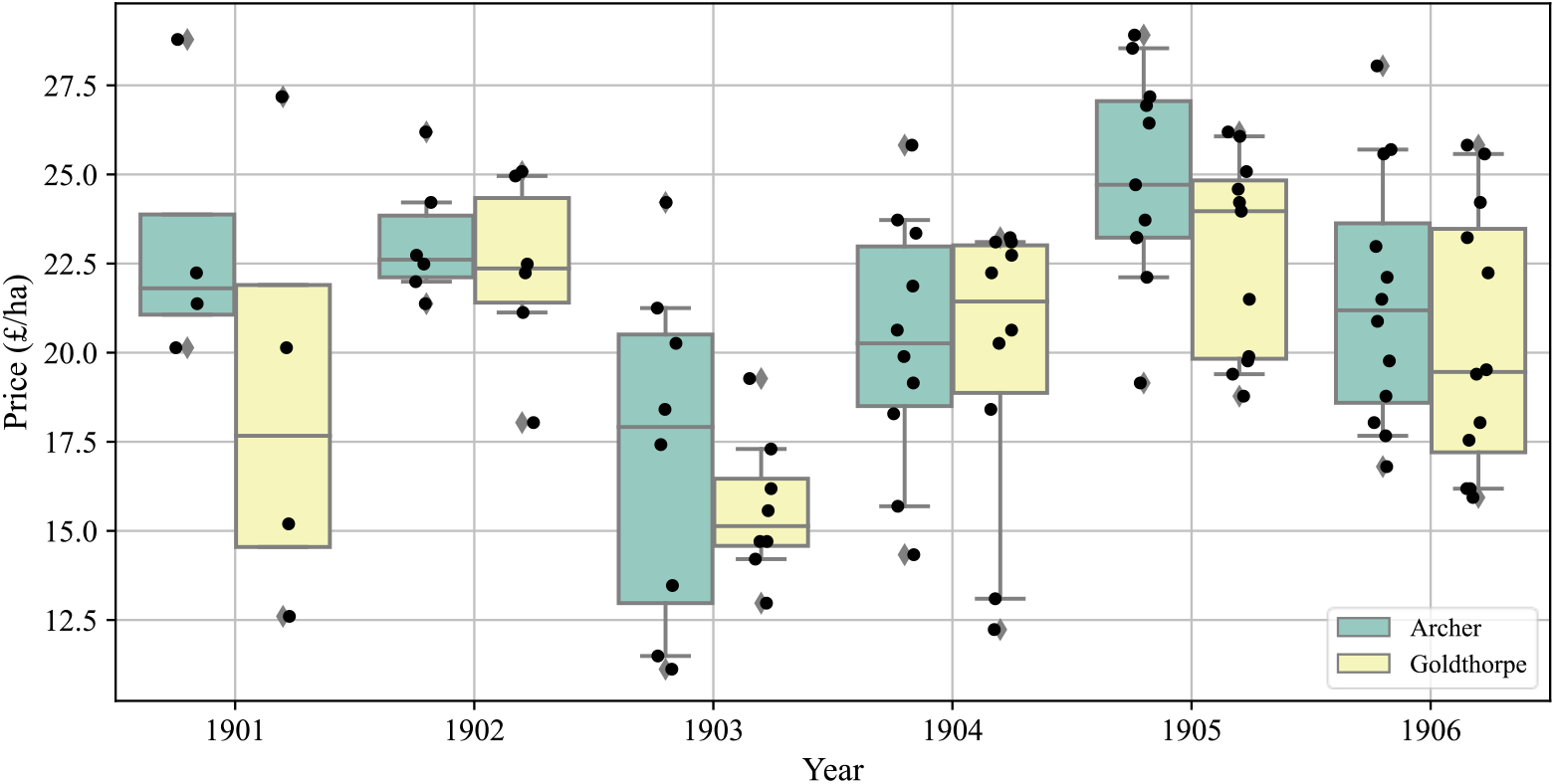
Barley trial price per hectare. (£/ha) (black dots) for 1901-1906 for two varieties: Archer (blue) and Goldthorpe (yellow). Outliers (diamonds) represent trial yields in the 5^th^ and 95^th^ percentiles. There were 51 trials per variety, increasing from 4 in 1901 to 12 in 1906. Data from Student (1923).

### 3.3 Irish climate analysis

#### 3.3.1 Long-term climate reveals anomalous years

Growing season rainfall anomalies show large interannual variability, with differences of up to 300 mm between neighbouring years (Figure 5). Averaged across all stations, the lowest yielding year 1903 was the wettest of the 6 barley trial years 1901-1906, with a large positive anomaly relative to the 1851-2010 average. Nationally the 1903 growing season received over 20% more rainfall than average. 1901 and 1902 were drier than average across the stations. Given the growing season in Ireland consistently receives high rainfall, water deficits are unlikely to be a yield-limiting factor.

**Figure 5:**
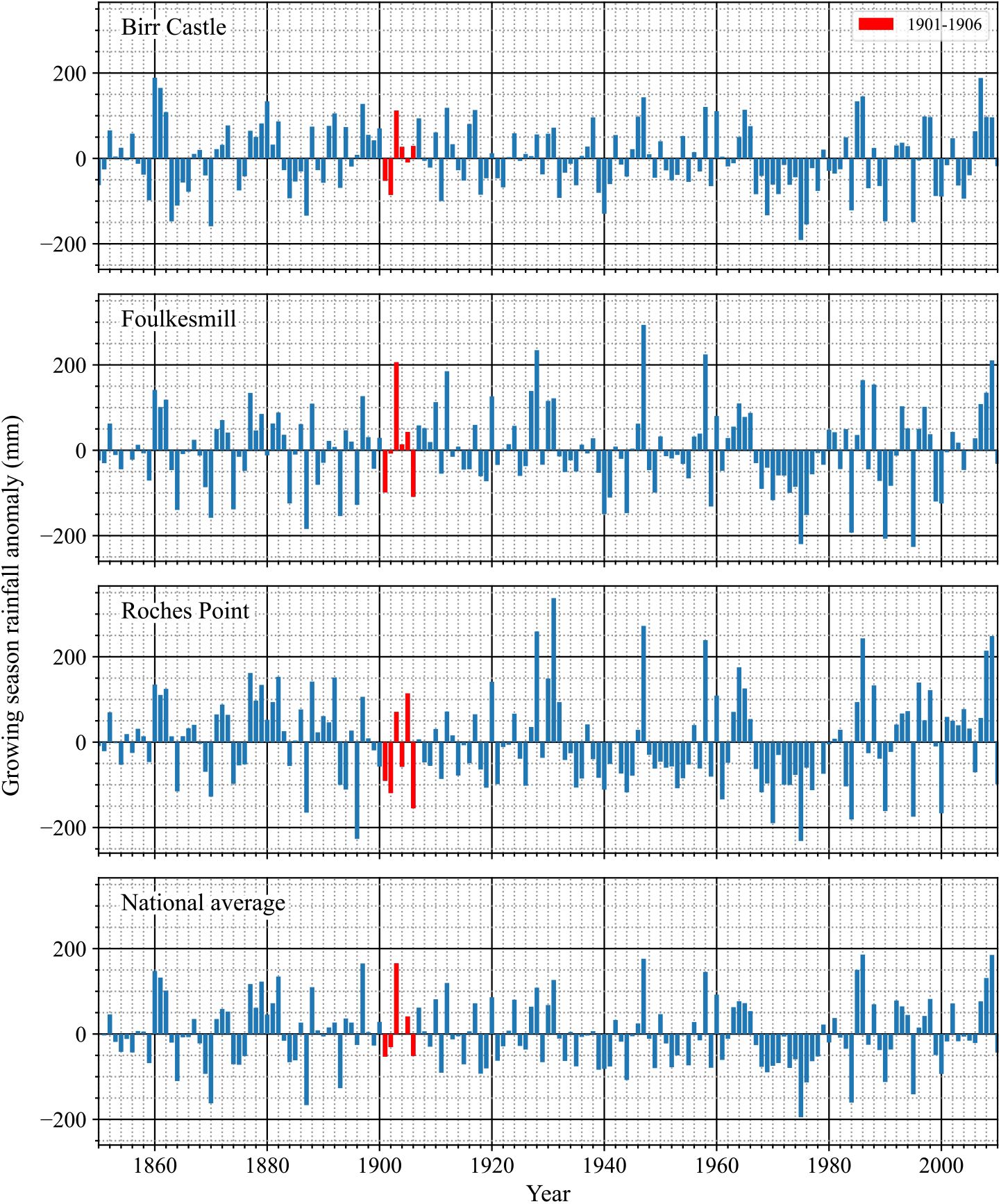
Growing season rainfall anomalies. (mm) (March-August) for Birr Castle, Foulkesmill and Roches Point stations and nationally for 1850-2010. Years 1901-1906 are shown in red. Here the national average anomaly is calculated using the Island of Ireland precipitation series from 25 stations.

Over the 1874-2020 period, significant long-term increases in growing degree days of 0.76°C yr^-1^ (r = 0.26, p=0.003) and 2.3°C yr^-1^ (r=0.66, p=3e-19) have been seen at Birr Castle and Glasnevin respectively (Figure 6). The more extreme increase in GDD seen at Dublin is likely due to increased urbanisation and industrialisation in the city (Dublin City Council 2017), decreasing the city’s albedo, increasing absorption of solar radiation and local temperatures. In addition to being the wettest of the 6 years, 1903 growing season has the 11^th^ lowest GDD recorded at both Birr Castle and Glasnevin stations across the period.

**Figure 6:**
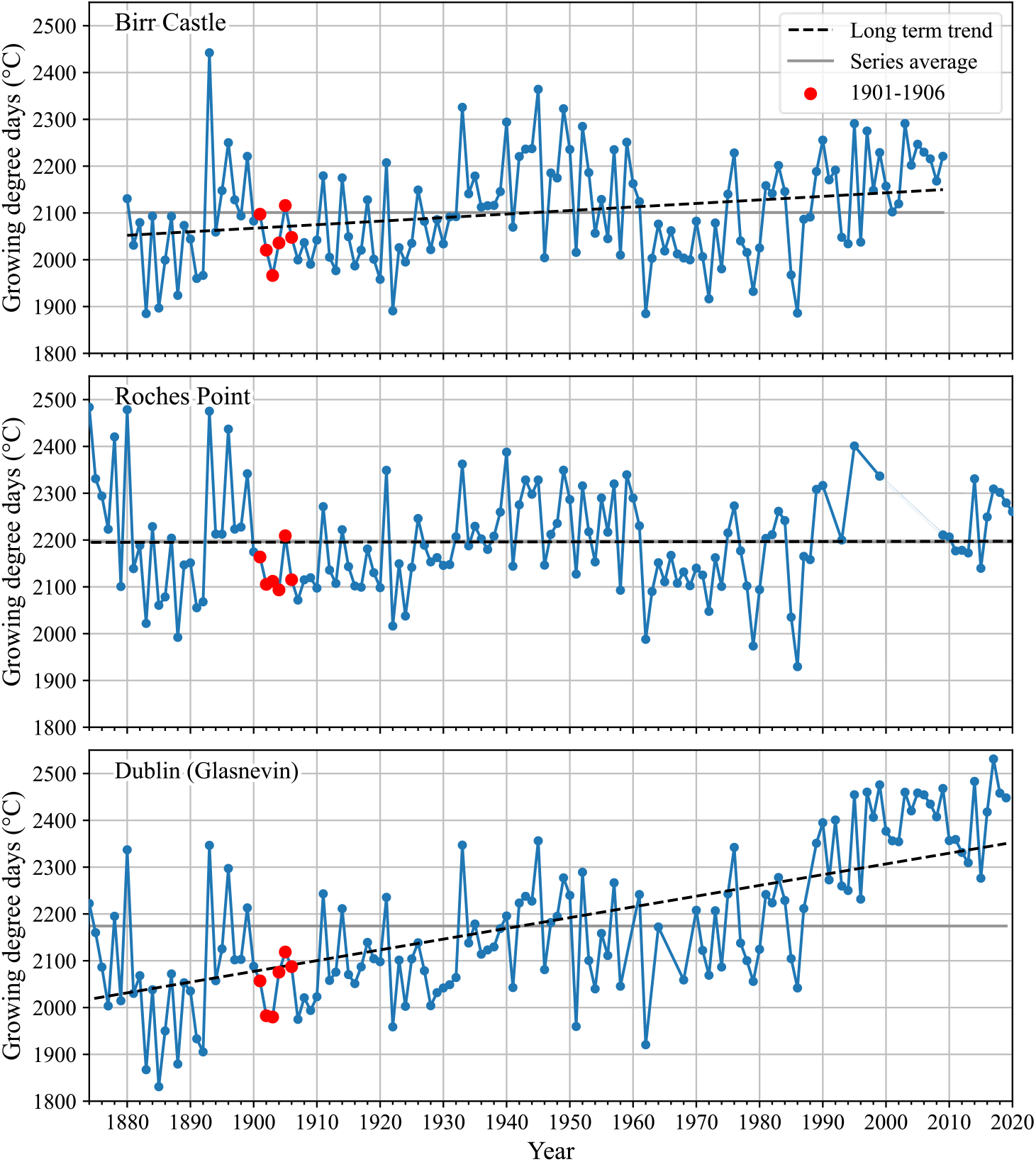
Growing season growing degree days. (°C) for Birr Castle, Roches Point and Glasnevin stations for 1874-2020. Growing degree days is the sum of the mean temperature on days when mean temperature is above 0°C from March to August. Roches Point is missing data for 1998-2008.

#### 3.3.2 1891-1920 climatology reveals extreme wetness in 1903

Comparing years 1901-1906 to the climate of 1891-1920 places the data in the context of the general climate at the time. The 6-year period showed some extreme wetness and temperatures.

March 1903 was the wettest March in the 1891-1920 30-year period for Ardee, Birr and Foulkesmill stations and nationally (Figure 7). The coastal stations Roches Point and Greenore saw more ‘normal’ rainfall amounts, with the former recording its highest March rainfall for the 30-year period in 1905. Cumulatively, 1903 was the wettest growing season in the 30-year period at Foulkesmill and nationally, recording over 600 mm rainfall. It was also the wettest growing season in the period 1901-1906 at Ardee, Birr and Greenore.

**Figure 7:**
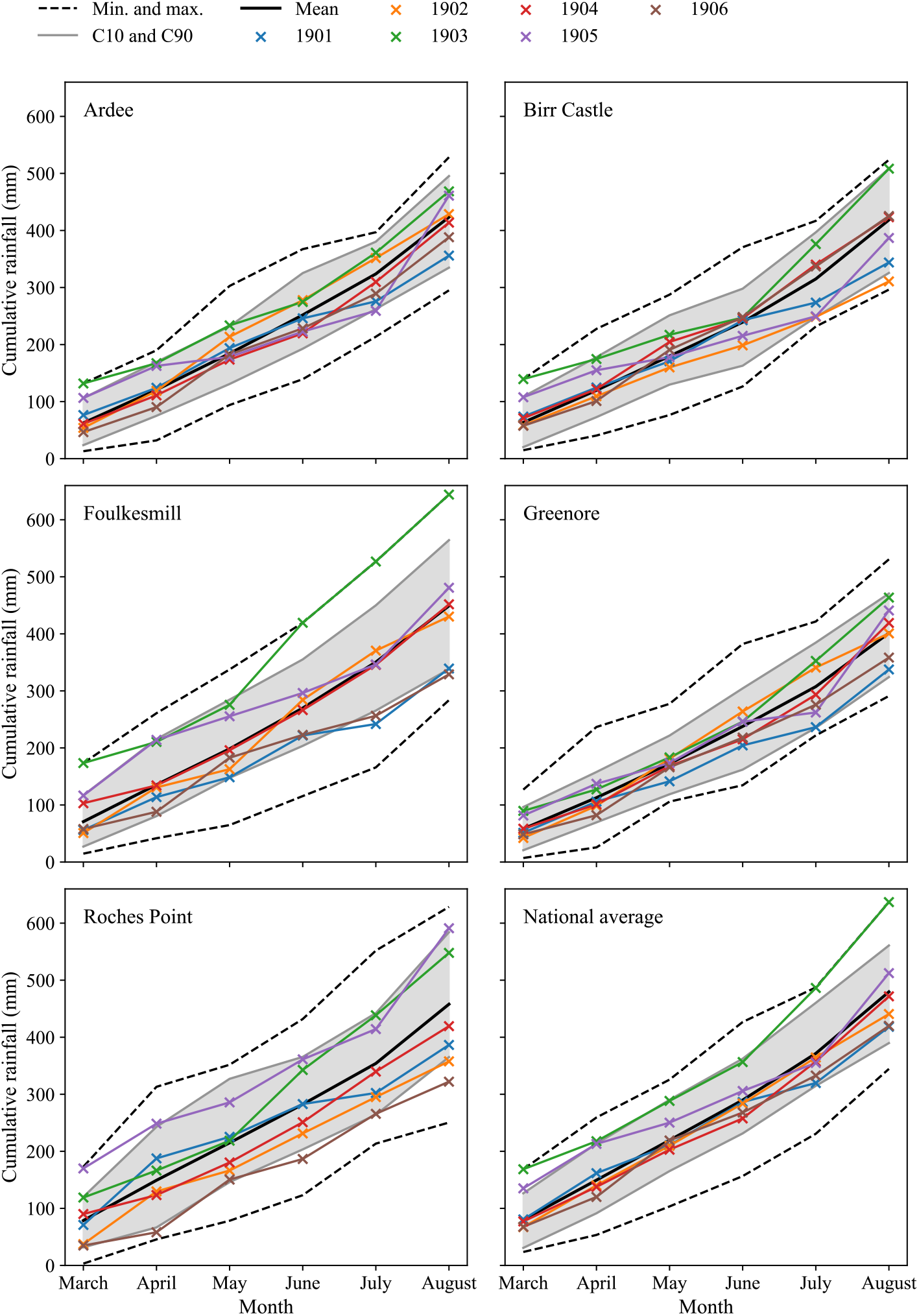
Cumulative monthly rainfall. (*mm) for Ardee, Birr Castle, Foulkesmill, Greenore, Roches Point stations and the national average across 25 stations for 1901, 1902, 1903, 1904, 1905 and 1906. The 1891-1920 average is shown (solid black line) along with the period 10^th^ and 90^th^ percentile values (grey lines) and the period minimum and maximum values (dashed black line). Note: Ardee station only has data for 1891-1913 therefore the averages are for this period instead. Data from* Ryan *et al*. (2020).

Roches Point’s coastal location is evident from the less extreme temperature values, with higher mean minimum temperatures and lower mean maximum temperatures (Figures 8 and 9). 1906 saw extremely low monthly mean minimum temperatures in April at all three stations, as well as the highest mean minimum temperature for August in the 30-years at Dublin. The mean minimum temperatures at Roches Point and Birr Castle for this month were closer to the average highlighting that climate extremes vary spatially and can be localised, contributing to the range in observed yields. March 1902 saw relatively high mean minimum and maximum temperatures whilst May 1902 saw much lower than average mean maximum temperatures (Figure 9).

**Figure 8:**
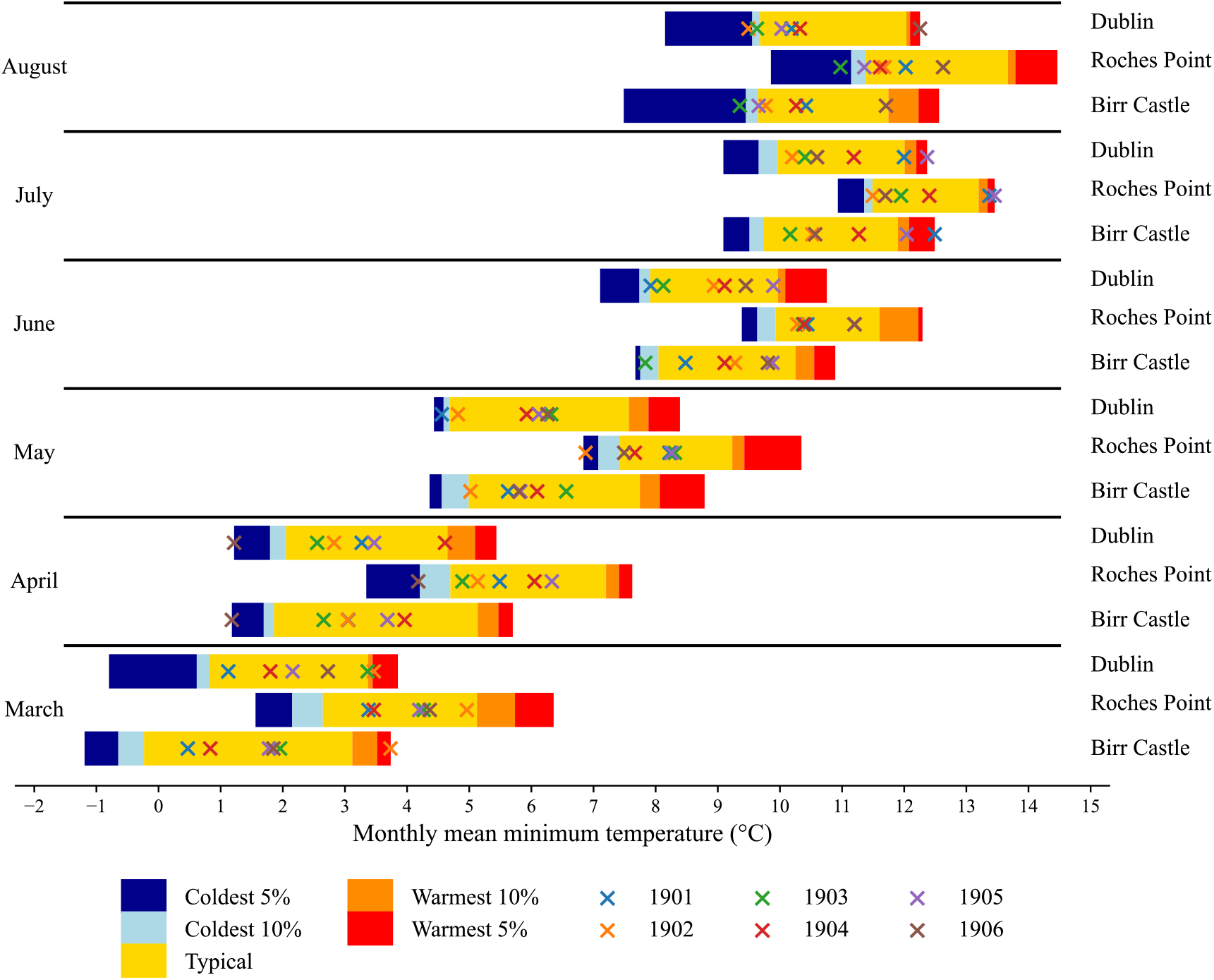
1891-1920 monthly mean minimum temperatures. (*°C) for Birr Castle, Roches Point and Dublin (Glasnevin) stations for the growing season. The range in temperatures for the coldest 5%, coldest 10%, warmest 10% and warmest 5% mean minimum temperatures are shown. Monthly mean minimum temperatures for 1901, 1902, 1903, 1904, 1905 and 1906 are also presented. Data from* (Mateus *et al*. 2020).

**Figure 9:**
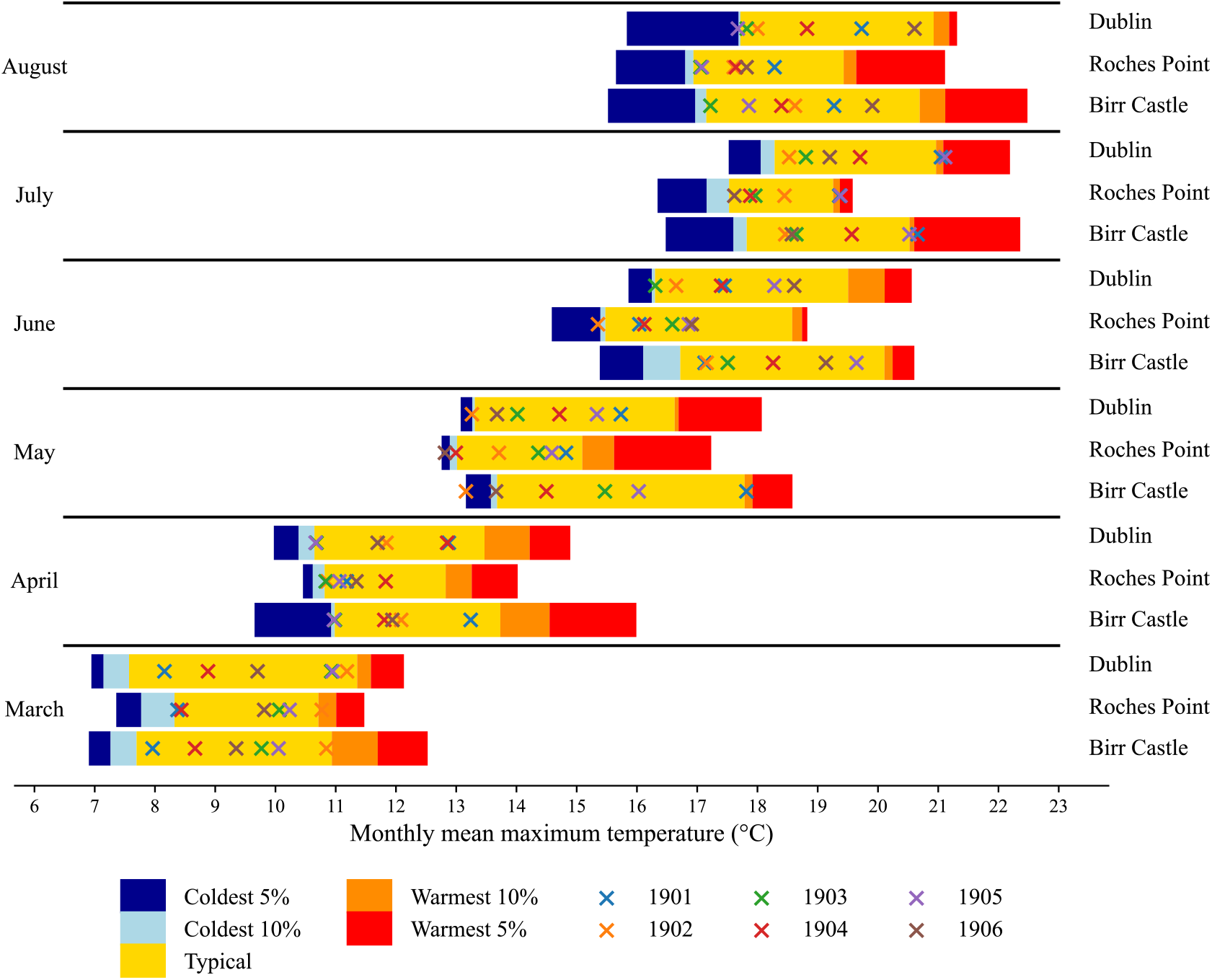
1891-1920 monthly mean maximum temperatures. (*°C) for Birr Castle, Roches Point and Dublin (Glasnevin) stations for the growing season. The range in temperatures for the coldest 5%, coldest 10%, warmest 10% and warmest 5% mean maximum temperatures are shown. Monthly mean maximum temperatures for 1901, 1902, 1903, 1904, 1905 and 1906 are also presented. Data from* (Mateus *et al*. 2020).

#### 3.3.3 Mapping climate anomalies

Analysis of growing season rainfall data from ERA-20C for 1901-1906 relative to the 1901-1930 averages shows that 1903 was much wetter than average across Ireland, the UK and much of Europe (Figure 10). 1906 was driest in the trials period. High rainfall is generally associated with a reduction in solar radiation and the 1903 growing season also received ~5% less PAR than the 1901-1930 average in Ireland (Figure 11). 1901 and 1904 show the most significant positive PAR anomalies over the growing season. Breaking this down into months, the 4 years 1901, 1902, 1904 and 1905 all have positive PAR anomalies in July in sync with the grain fill period.

**Figure 10:**
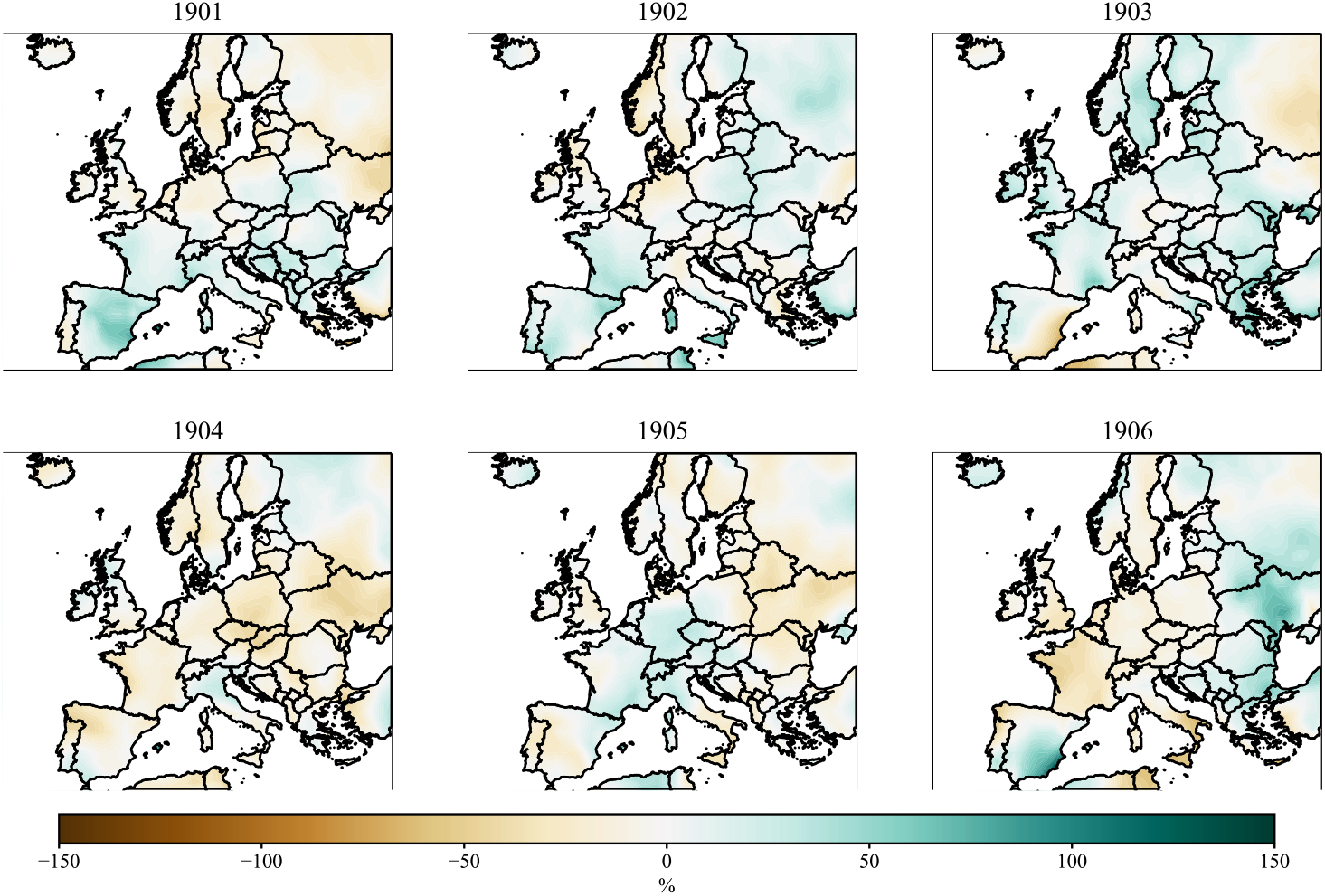
Growing season (March to August) rainfall anomalies. (*%) relative to the 1901-1930 average. Brown corresponds to drier than average and blue corresponds to wetter than average. Data from ERA-20C* (Poli *et al*. 2016).

**Figure 11:**
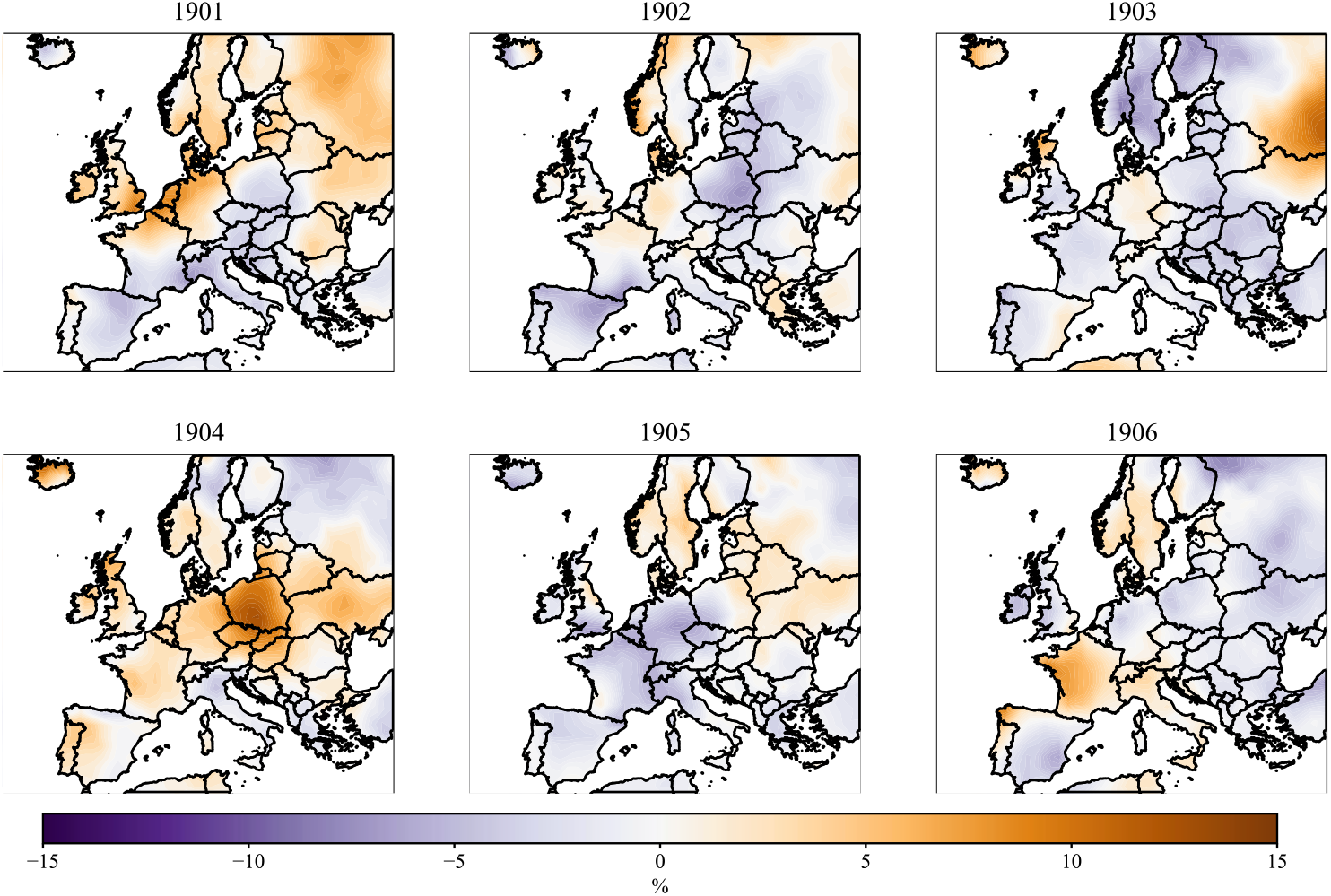
Growing season (March to August) total photosynthetically active radiation. (*PAR) anomalies (%) relative to 1901-1930 average. Purple corresponds to less PAR than average whilst orange corresponds to more PAR than average. Data from ERA-20C (Poli et al. 2016*).

1905 was the only growing season in the period when Ireland had a positive temperature anomaly, of ~0.3°C. Higher than average temperatures were also experienced across the UK and most of central, eastern and northern Europe (Figure 12). 1903 was the coldest growing season in Ireland, about 1°C below the 1901-1930 average.

**Figure 12:**
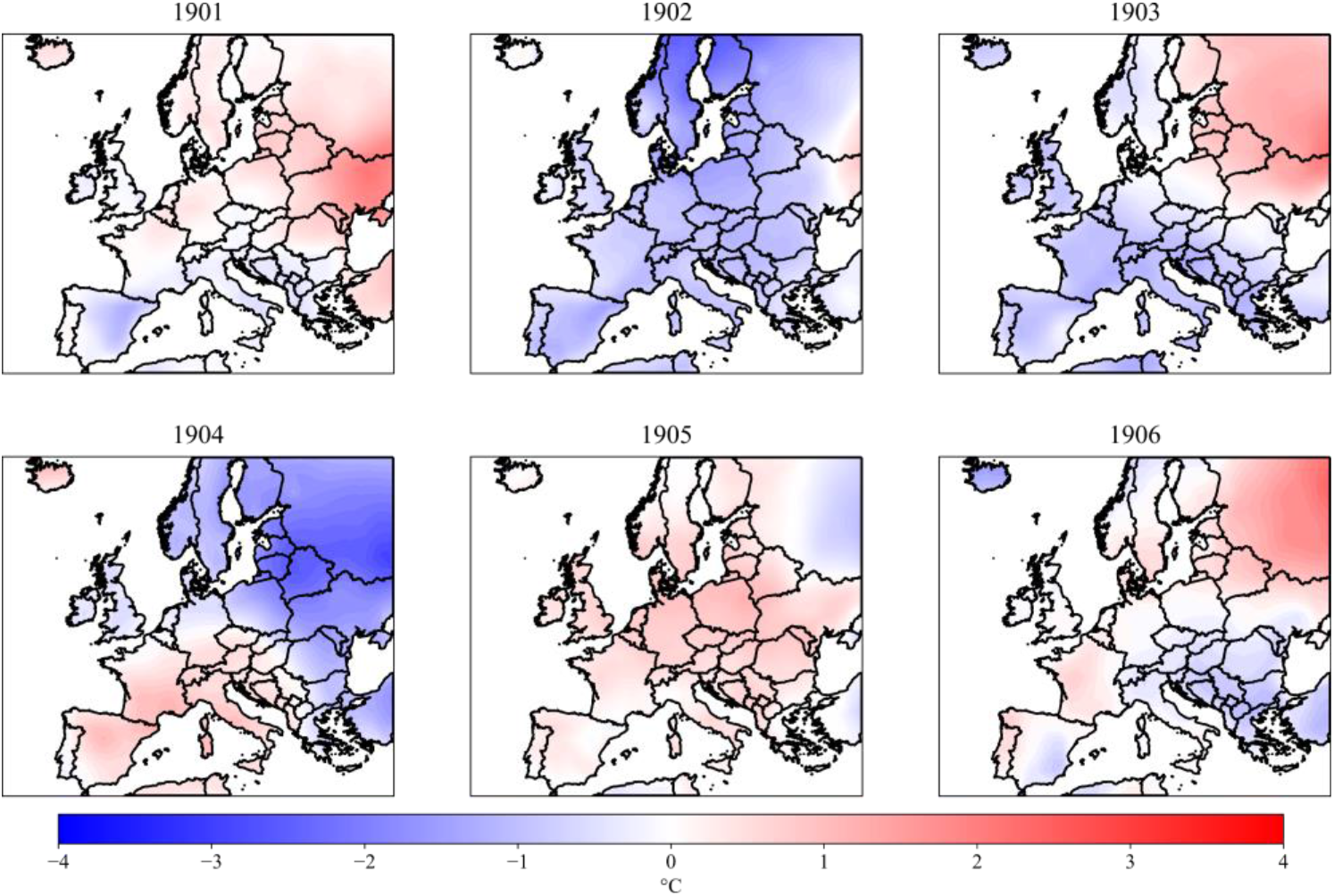
Growing season (March to August) mean temperature anomalies. (*°C) relative to 1901-1930 average. Blue corresponds to colder than average whilst red corresponds to warmer than average. Data from ERA-20C* (Poli *et al*. 2016).

### 3.4 Modelling Irish spring barley and climate

#### 3.4.1 Variable selection methods

We found the best subset selection, forwards and backwards selection and the elastic net methods failed to sufficiently simplify the climate model to then include the selected climate covariates in the mixed model with year, variety and site [2] (Table S2). A combination of using too many highly correlated variables (Figure S1) and too few farm growing seasons likely contributed to this. The worst performing was backwards stepwise selection which did not drop any variables. The two lasso methods reduced the model complexity significantly from 19 to less than 7 climate variables, but these models had very low adjusted R^2^ values of close to 0, indicating a poor model fit (Table S3). Using the mixed-model backwards elimination approach, we found that all the climate variables were dropped. In all methods except this, July maximum temperature was kept in the model.

#### 3.4.2 Principal Component Analysis

Using PCA we found that the first 6 principal components (PC) explained ~90% of the observed variation in yield (Table S4). These were then input into equation [3], replacing the climate variables. PCs 2, 3, 4 and 5 were significant. However, the 4 PCs were not clearly defined by just one or two climate variables, rather several. Using PCA didn’t therefore simplify the model.

#### 3.4.3 Pearson’s correlation analysis

In yield-climate correlation analysis we found July rainfall and July maximum temperature have the largest absolute correlation with yield: −0.49 and 0.45, respectively (Figure S2). These variables have a strong negative correlation.

#### 3.4.4 Akaike Information Criterion

To understand if adding temperature or rainfall climate variables to the mixed model [3] improves the fit, the AIC of the mixed model of [3] without any climate covariates was first calculated (Table S5). We then added each climate variable and its interaction with variety to the model one at a time. None of the interactions with variety were significant, the variety x climate interaction term was dropped from the model and the models with each climate variable were looped through again.

Only three variables – July maximum temperature, August maximum temperature and July total rainfall – were significant when included in the model. The models which included either July maximum temperature or August maximum temperature improved the AIC and model fit. Notably all the models that contained temperature had a lower AIC and better fit than any of the rainfall models, including the significant July rainfall model (Table S5).

Both July mean maximum temperature and August mean maximum temperature had a positive relationship with yield (Table S5), such that yield increased by ~ 1/4 t/ha per 1°C increase in July mean maximum temperature and by ~ 1/5 t/ha per 1°C increase in August maximum temperature.

### 3.5 Comparison of standard error of difference between means

The mean difference in the variety values is £1.52/ha (12 shillings/acre) with a standard deviation of £2.95/ha (23.9 shillings/acre) and corresponding standard error of the mean difference £0.41/ha (3.3 shillings/acre), in accordance with Student (1923). This corresponds to a t-statistic of 3.680, which was statistically significant (p=0.0006) at the 95% level (df = 50). This provided strong evidence that there was a difference in varietal performance.

Calculating the standard error of difference between variety values in the three mixed models containing significant climate effects (Table S5) gives identical values (to 2 s.f.) of £0.41/ha (3.3 shillings/acre). This indicates that this model does not reduce the standard error, which is expected given no variety x climate interactions were included in the final models. The climate variables simply partitioned the effects of Farm and Year and did not affect the Variety effect.

## 4 Discussion

### 4.1 Climatic causes of yield variability

Use of recently digitised weather data for the early 20^th^ century has allowed us to show that contrasting climatic conditions in 1903 and 1905 coincided with variation in spring barley varietal performance. In 1903 a wet March (Figure 7) likely made it challenging to drill the crop, resulting in delayed planting shortening the growing season and compounded by potential difficulties in crop establishment. Nationally, the 1903 growing season received over 20% more rainfall than average (Figure 5), contributing to greater cloud coverage and lower than average growing season PAR (Figure 11), notably during April, May, June and July. Reduced solar radiation interception during the grain fill period in June and July constrains photosynthesis, reducing the final ear weight amassed in this period (TEAGASC 2017). The 1903 growing season was also cooler than average (Figure 12) with low GDDs (Figure 6). This coincides with the year of lowest mean yields and greatest yield variability for *Archer*, but much lower variability for *Goldthorpe* (SD = 0.22 t/ha) (Figure 2).

A more recent experiment detailed by Gothard *et al*. (1983) found that *Goldthorpe* outperformed *Archer* when spring and summer rainfall was high. Along with our results, this suggests *Goldthorpe* may be able to withstand much higher soil moisture and waterlogging. Hunter (1929) notes that *Goldthorpe* requires plenty of moisture to produce the best yields and quality, supporting this theory. Continual dampness can also increase disease pressures for diseases such as Barley Scald (*Rhynchosporium*) which prefer cool wet weather and which, if present early in the season, can reduce tiller survival and potential yields (TEAGASC 2017). If Barley Scald was present, this may show greater resistance of *Goldthorpe* to the disease.

In contrast, the 1905 growing season was warmer than average (Figure 12) with high growing season GDDs (Figure 6). There was low growing season PAR (Figure 11), but high PAR in July, when high solar radiation is important for grain fill. The growing season was drier than average, starting wet but drying in June and July. These favourable conditions likely contributed to the relatively high yields seen in 1905 for both varieties.

Of the farms with 6 years of trials data, Farmer Wolfe performed the best on average (Figure 3). This farm was located ~30km south-west of Birr Castle and experienced higher summer temperatures and less summer rainfall than the other two farms. Other factors such as favourable agronomy, farm management and soil type may also have encouraged higher yields here.

### 4.2 Statistical methods

Through trialling various variable selection methods, we have highlighted the importance of identifying collinearity early on in analysis involving multiple covariates. The use of these methods and PCA was limited by the high correlation between covariates within a small dataset, but it was still possible to extract information on the most important variables using simple mixed models. We were able to show that July maximum temperature and August maximum temperature had a positive relationship with yield and that July total rainfall had a negative relationship with yield (Table S5). July rainfall can also be used as a proxy for solar radiation, so a wet July would usually be associated with more cloud cover, reducing solar radiation interception during grain fill. Likewise wet weather during grain filling can encourage ear and grain diseases, such as fusarium ear blight and ergot, which can cause shrivelled grain and mycotoxins (AHDB 2018). Hence the plant benefits from more solar radiation and less rainfall in July. Higher July maximum temperature implies less daytime cloud cover intercepting solar radiation, hence the correlation between these two July variables and yield is of opposite polarity. In future analysis of more recent crop yield data, inclusion of solar radiation data in the models would be desirable to directly quantify the relationship between solar radiation and yield.

Our finding that July temperatures are positively correlated with spring barley yield contrasts with other published research which shows that warmer temperatures during anthesis and grain fill can have a detrimental effect (Addy *et al*. 2021; Hakala *et al*. 2020). This result is highly likely due to July maximum temperatures in Ireland in the early 20^th^ century falling well short of those more regularly seen today in some major UK spring barley growing areas. Specifically, maximum temperature did not exceed 28°C during the 6-year trials period whereas those in South-East England now regularly exceed 30°C in summer months. This finding shows the importance of region-specific crop-climate research: despite the proximity of the UK to Ireland their climates differ and the same relationships between weather variables and yield cannot be assumed.

We were unable to detect any GxE within the mixed models used. The lack of significance throughout of climate variety interactions may well be related to the relatively small trials dataset, approximation of site locations and sometimes large distances to weather stations. However, it is clear from the higher performance of *Goldthorpe* in 1903 relative to *Archer* coupled with wider evidence (Gothard *et al*. 1983; Reid *et al*. 1929) that GxE is a driver of performance here. This highlights the importance of considering the local climate in crop variety selection.

The last few years have seen a surge in the growing of heritage barley varieties from the early 20^th^ century. *Goldthorpe*, its predecessor *Chevalier* and offspring *Irish Goldthorpe*, as well as hybrids of *Archer*, such as *Plumage Archer*, have been grown for breweries across the UK and Ireland and are currently being investigated by organisations such as New Heritage Barley. Some heritage varieties display highly desirable traits, such as Fusarium fungal disease resistance in *Chevalier* (BBSRC UKRI 2016). How these varieties perform in the current and future climate is of interest given the performance of these varieties in the 1901-1906 trials. It is hoped that *Archer* and *Goldthorpe* will be trialled on large scale field plots to allow for comparisons with the yields from 1901-1906, but also to test the models in the current climate on larger datasets.

### 4.3 Historical perspective of Student’s 1923 paper

Student’s comments in 1923 on the requirement for large scale farm testing remains as relevant today. In recent years a greater emphasis is also placed on grower input through participatory breeding approaches (Ceccarelli *et al*. 2007; Weltzien & Christinck 2017) and large-scale farm trials in strip tests (Lacoste *et al*. 2022; Marchant *et al*. 2019; Piepho *et al*. 2011).

Student states that the advantage for the farmer of large scale trials is that s/he “.. .always has a healthy contempt for gardening” and … “some varieties which have come out well on the small scale have not done as well in the field, though this is not at all common”. That said, two-acre plots (0.4 ha) are very large, as Student recognises, even for large-scale plots, though the produce was also intended to provide seed for subsequent manufacturing tests, presumably including malting though we have no record that they took place.

The importance of collaboration is also commented on: here between farmers in carrying out large scale experiments …”. it is only by co-operation [between farmers] that enough evidence can be obtained to be of any value”, though he sees such co-operation as being most likely co-ordinated by government bodies. It is unfortunate that, to date, that hasn’t happened.

Another laudable feature of Student’s paper is that he made the data available. Admittingly this was largely to illustrate the method of analysis, but full data release is still not the norm. Subsequently, the data was reanalysed by Patterson (1997), also for educational purposes. We do not know whether Student had soil and weather records available to him (he was analysing the yield data when it was already twenty years old) or whether he would have felt it advantageous to include them. In fact, we find near identical results to Student: *Archer* yields more than *Goldthorpe*. In the absence of any detectable variety x climate variable interactions (as here), this is expected. The climatic variables which are available to us have, however, been used to identify drivers of yield differences between sites and years in a dataset ~120 years old.

## 5 Conclusion

Through combining recently published historical rainfall and temperature data with spring barley trials data, it has been possible to identify climatic influences on spring barley yield variability seen in early twentieth century trials data in Ireland, building on the earlier findings of Student (1923).

Despite being available for ~100 years, we have demonstrated that there is value in adding historical climate data to this small trials dataset. Today’s large-scale trial datasets provide a great opportunity for further insight on crop-climate interactions in a changing climate.

## Acknowledgements

The authors are grateful for the help provided by Ciara Ryan and Mary Curley at Met Éireann in providing access to the climate data, in particular the daily rainfall data which was unpublished at the time. The authors have no competing interests to declare.

## References

Addy, J. W. G., Ellis, R. H., Macdonald, A. J., Semenov, M. A., & Mead, A. (2021). Changes in agricultural climate in South-Eastern England from 1892 to 2016 and differences in cereal and permanent grassland yield. Agricultural and Forest Meteorology, 308–309(July). doi:10.1016/j.agrformet.2021.108560

AHDB. (2018). Wheat and barley disease management guide.

Baron, V. S., Aasen, A., Oba, M., … Stevenson, C. F. (2012). Swath-grazing potential for small-grain species with a delayed planting date. Agronomy Journal, 104(2), 393–404.

Bates, D., Mächler, M., Bolker, B. M., & Walker, S. C. (2020). lme4: linear mixed-effects models. R package version 1.1.21. Retrieved from https://github.com/lme4/lme4/

Bauer, A., Frank, A. B., & Black, A. L. (1992). A crop calendar for spring wheat and for spring barley. North Dakota Farm Research, 49(6), 21–25.

BBSRC UKRI. (2016). New Heritage Barley Ltd – reviving a Victorian barley variety for modern brewers. Retrieved from https://bbsrc.ukri.org/documents/1606-new-heritage-barley-pdf/

Bindereif, S. G., Rüll, F., Kolb, P., … Gebauer, G. (2021). Impact of global climate change on the european barley market requires novel multi-method approaches to preserve crop quality and authenticity. Foods, 10(7). doi:10.3390/foods10071592

Ceccarelli, S., Grando, S., & Baum, M. (2007). Participatory plant breeding in water-limited environments. Experimental Agriculture, 43(4), 411–435.

de los Campos, G., Pérez-Rodríguez, P., Bogard, M., Gouache, D., & Crossa, J. (2020). A data-driven simulation platform to predict cultivars’ performances under uncertain weather conditions. Nature Communications, 11(1). doi:10.1038/s41467-020-18480-y

Dublin City Council. (2017). Dublin City Parks Strategy 2017-2022. Retrieved from https://www.dublincity.ie/dublin-city-parks-strategy/2-parks-and-landscapes-perspective/24-international-perspective/241-population-and-urban-areas

Fabio, E. S., Kemanian, A. R., Montes, F., Miller, R. O., & Smart, L. B. (2017). A mixed model approach for evaluating yield improvements in interspecific hybrids of shrub willow, a dedicated bioenergy crop. Industrial Crops and Products, 96, 57–70.

Gillberg, J., Marttinen, P., Mamitsuka, H., & Kaski, S. (2019). Modelling G×E with historical weather information improves genomic prediction in new environments. Bioinformatics, 35(20), 4045–4052.

Gothard, P. G., Riggs, T. J., & Smith, D. B. (1983). The Malting Quality of Some Spring Barley Varieties Grown in England and Wales Between 1880 and 1980. Journal of the Institute of Brewing, 89(5), 344–348.

Hakala, K., Jauhiainen, L., Rajala, A. A., Jalli, M., Kujala, M., & Laine, A. (2020). Different responses to weather events may change the cultivation balance of spring barley and oats in the future. Field Crops Research, 259. doi:10.1016/j.fcr.2020.107956

Hunter, H. (1913). Irish Barley Growing Experiments. Journal of the Institute of Brewing, 547–577.

Ivandic, V., Hackett, C. A., Zhang, Z. J., … Forster, B. P. (2000). Phenotypic responses of wild barley to experimentally imposed water stress. Journal of Experimental Botany, 51(353), 2021–2029.

Juskiw, P. E., Jame, Y. W., & Kryzanowski, L. (2001). Phenological development of spring barley in a short-season growing area. Agronomy Journal, 93(2), 370–379.

Kahiluoto, H., Kaseva, J., Balek, J., … Trnka, M. (2019). Decline in climate resilience of European wheat. Proceedings of the National Academy of Sciences, 116(1), 123–128.

Kuznetsova, A., Brockhoff, P. B., & Christensen, R. H. B. (2017). lmerTest Package: Tests in Linear Mixed Effects Models. Journal of Statistical Software, 82(13). doi:10.18637/jss.v082.i13

Lacoste, M., Cook, S., McNee, M., … Hall, A. (2022). On-Farm Experimentation to transform global agriculture. Nature Food, 3(1), 11–18.

Lopes, M. S. (2022). Will temperature and rainfall changes prevent yield progress in Europe? Food and Energy Security, (March 2021), 1–12.

Malcolm, J. P. (1983). The History of Barley in New Zealand. Agronomy Society of NZ Special Publication, 3–15.

Marchant, B., Rudolph, S., Roques, S., … Sylvester-Bradley, R. (2019). Establishing the precision and robustness of farmers’ crop experiments. Field Crops Research, 230(November 2017), 31–45.

Mateus, C., Potito, A., & Curley, M. (2020). Reconstruction of a long-term historical daily maximum and minimum air temperature network dataset for Ireland (1831-1968). Geoscience Data Journal, 7(2), 102–115.

Murphy, C., Broderick, C., Burt, T. P., … Wilby, R. L. (2018). A 305-year continuous monthly rainfall series for the island of Ireland (1711-2016). Climate of the Past, 14(3), 413–440.

Patterson, H. D. (1997). Analysis of series of variety trials. In Statistical methods for plant variety evaluation, Dordrecht: Springer, pp. 139–161.

Piepho, H. P., Richter, C., Spilke, J., Hartung, K., Kunick, A., & Thöle, H. (2011). Statistical aspects of on-farm experimentation. Crop and Pasture Science, 62(9), 721–735.

Poli, P., Hersbach, H., Dee, D. P., … Fisher, M. (2016). ERA-20C: An atmospheric reanalysis of the twentieth century. Journal of Climate, 29(11), 4083–4097.

Reid, R., Hunter, H., Stewart, J., … Keen, B. A. (1929). Malting Barley. Rothamsted Conferences, London.

Rezaei, E. E., Siebert, S., & Ewert, F. (2015). Intensity of heat stress in winter wheat - Phenology compensates for the adverse effect of global warming. Environmental Research Letters, 10(2). doi:10.1088/1748-9326/10/2/024012

Ryan, C., Murphy, C., McGovern, R., Curley, M., & Walsh, S. (2020). Ireland’s pre-1940 daily rainfall records. Geoscience Data Journal. doi:10.1002/gdj3.103

Schloerke, B., Cook, D., Larmarange, J., … Crowley, J. (2021). GGally: Extension to “ggplot2”. Version 2.1.2. Retrieved from https://cran.r-project.org/package=GGally

Schmidt, S. B., George, T. S., Brown, L. K., … Husted, S. (2019). Ancient barley landraces adapted to marginal soils demonstrate exceptional tolerance to manganese limitation. Annals of Botany, 123(5), 831–843.

Student. (1923). On Testing Varieties of Cereals. Biometrika, 15(3/4), 271–293.

TEAGASC. (2017). The Spring Barley Guide. Crops Environment & Land Use Programme. Retrieved from https://www.teagasc.ie/media/website/publications/2015/The-Spring-Barley-Guide.pdf

TEAGASC. (2020). Spring Cereals. Retrieved October 26, 2020, from https://www.teagasc.ie/crops/crops/cereal-crops/spring-cereals/

Tibshirani, R. (1996). Regression Shrinkage and Selection Via the Lasso. Journal of the Royal Statistical Society: Series B (Methodological), 58(1), 267–288.

Trnka, M., Eitzinger, J., Dubrovský, M., … Žalud, Z. (2010). Is rainfed crop production in central Europe at risk? Using a regional climate model to produce high resolution agroclimatic information for decision makers. Journal of Agricultural Science, 148(6), 639–656.

Weltzien, E., & Christinck, A. (2017). Participatory Breeding: Developing Improved and Relevant Crop Varieties With Farmers. In S. Snapp & B. Pound, eds., Agricultural Systems: Agroecology and Rural Innovation for Development, 2nd edn, Elsevier Inc., pp. 259–301.

Wu, X., Cai, K., Zhang, G., & Zeng, F. (2017). Metabolite Profiling of Barley Grains Subjected to Water Stress: To Explain the Genotypic Difference in Drought-Induced Impacts on Malting Quality. Frontiers in Plant Science, 8(September), 1–12.

Zou, H., & Hastie, T. (2005). Regularization and variable selection via the elastic net. Journal of the Royal Statistical Society. Series B: Statistical Methodology, 67(2), 301–320.

