## Supplementary Tables and Figures for "Combining historical agricultural and climate datasets sheds new light on early 20^th^ century barley performance"

**Figure S1: Correlation between each mean maximum and minimum monthly temperature and total rainfall variables.** *Only significant correlations (p < 0.05) are shown. The larger the square the stronger the correlation. Dark red corresponds to strong negative correlations, dark blue corresponds to strong positive correlations.*

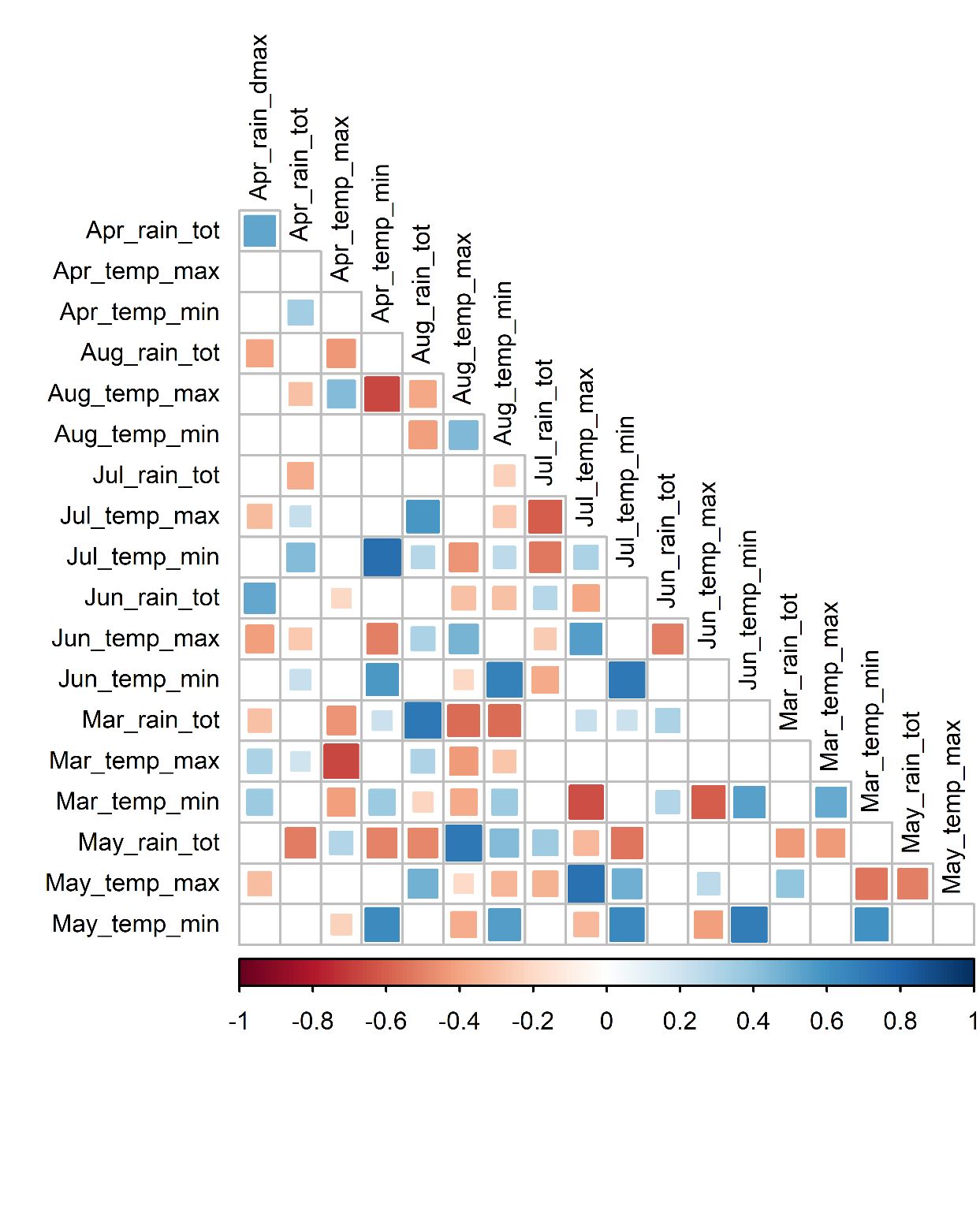

**Figure S2: Correlation between each monthly climate variable and yield.**
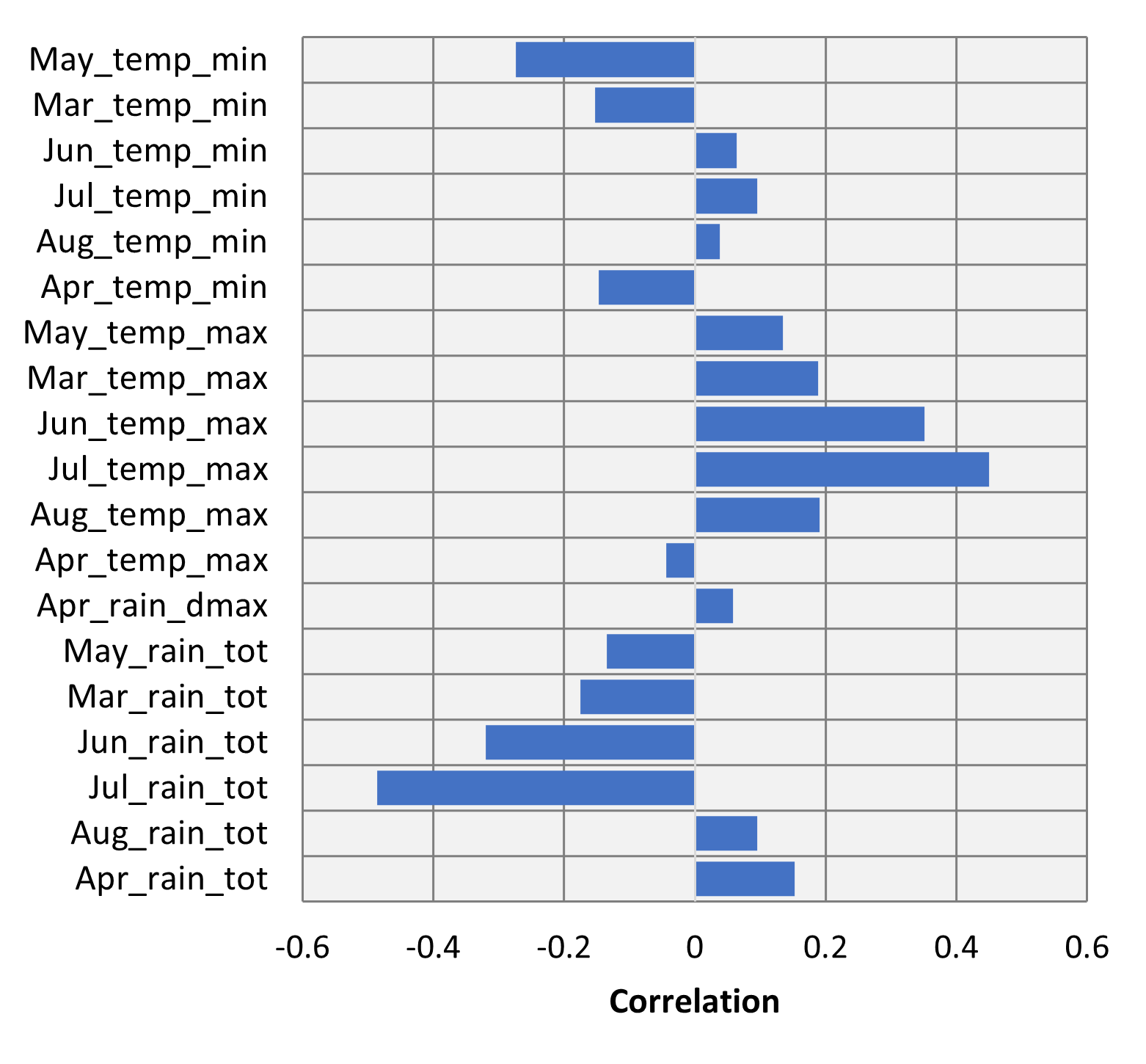

Tables

**Table S1: Methods of variable selection used to select climate covariates**

|  | **Method** | **Functions** | **Packages** | **Arguments** | **Reference** |
| --- | --- | --- | --- | --- | --- |
| 1 *= cv.b* | Best subset selection using cross-validation | *regsubsets* | **tidyverse**  **caret**  **leaps** | nvmax = 19 | (Wickham *et al.* 2019)  (Kuhn 2020)  (Lumley 2020) |
| 2a = *s.bo* | Stepwise selection in both directions | *stepAIC* | **MASS** | direction = “both” | (Venables & Ripley 2002) |
| 2b = *s.ba* | Backwards stepwise selection | *trainControl* | **leaps**  **caret**  **tidyverse** | number = 10 | (Kuhn 2020)  (Lumley 2020)  (Wickham *et al.* 2019) |
|  |  | *train* |  | method = “leapBackward” |  |
| 2c *= s.f* | Forwards stepwise selection | *trainControl* | **leaps**  **caret**  **tidyverse** | number = 10 | (Kuhn 2020)  (Lumley 2020)  (Wickham *et al.* 2019) |
|  |  | *train* |  | method = “leapForward” |  |
| 3a = *l.m* | Lasso using optimal λ that minimises the cross-validation error | *cv.glmnet* | **tidyverse**  **caret**  **glmnet** | family = “gaussian”, alpha = 1 | (Wickham *et al.* 2019)  (Lumley 2020)  (Friedman *et al.* 2010) |
|  |  | *glmnet* |  | family = “gaussian”, alpha = 1  lambda = cv.lasso$lambda.min |  |
| 3b = *l.1* | Lasso using λ with gives the simplest model and lies within one standard error of lambda.min | *cv.glmnet* | **tidyverse**  **caret**  **glmnet** | family = “gaussian”, alpha = 1 | (Wickham *et al.* 2019)  (Lumley 2020)  (Friedman *et al.* 2010) |
|  |  | *glmnet* |  | family = “gaussian”, alpha = 1  lambda = cv.lasso$lambda.1se |  |
| 4 = *e.n* | Elastic net using optimal λ and that minimise the cross-validation error | *trainControl* | **tidyverse**  **caret**  **glmnet** | method = “repeatedcv”, number = 10, repeats = 5 | (Wickham *et al.* 2019)  (Lumley 2020)  (Friedman *et al.* 2010) |
|  |  | *train* |  |  |  |
|  |  | *glmnet* |  | method = “glmnet”,  tuneLength = 10 |  |
|  |  |  |  | family = “gaussian” |  |

| **Variable** | **Coefficients** | | | | | | | | **p-value** | | | | | **sig. (p < 0.05)** | | | |
| --- | --- | --- | --- | --- | --- | --- | --- | --- | --- | --- | --- | --- | --- | --- | --- | --- | --- |
|  | c.m | cv.b | s.bo | s.ba | s.f | l.m | l.1 | e.n | c.m | s.bo | s.ba | s.f | c.m | | s.bo | s.ba | s.f |
| (Intercept) | 2.798 | 2.798 | 2.798 | 2.798 | 2.798 | 2.815 | 2.799 | 2.822 |  |  |  |  |  | |  |  |  |
| apr_rain_dmax | 0.423 | 0.503 | 0.348 | 0.423 | 0.458 | 0.073 | 0 | 0.132 | 0.0002 | 0.3096 | 0.0002 | 0.0021 | * | |  | * | * |
| apr_rain_tot | -0.241 | -0.356 | -0.212 | -0.241 | -0.345 | 0 | 0 | -0.023 | 0.0423 | 0.0403 | 0.0423 | 0.0432 | * | | * | * | * |
| apr_temp_max | -0.380 | -0.373 | -0.190 | -0.380 | -0.298 | 0 | 0 | 0 | 0.0131 | 0.0921 | 0.0131 | 0.1208 | * | |  | * |  |
| apr_temp_min | 1.450 | 0.852 | 1.048 | 1.450 | 0.771 | 0 | 0 | 0.051 | 0.4955 | 0.0683 | 0.4955 | 0.1790 |  | |  |  |  |
| aug_rain_tot | 0.174 |  |  | 0.174 |  | 0 | 0 | -0.012 | 0.2541 |  | 0.2541 |  |  | |  |  |  |
| aug_temp_max | 0.392 | 0.722 |  | 0.392 | 0.603 | 0 | 0 | 0.082 | 0.0068 |  | 0.0068 | 0.9660 | * | |  | * |  |
| aug_temp_min | 0.732 |  | 0.762 | 0.732 | 0.125 | 0 | 0 | 0.096 | 0.0323 | 0.5125 | 0.0323 | 0.2499 | * | |  | * |  |
| jul_rain_tot | -0.169 |  |  | -0.169 | -0.017 | -0.154 | -0.095 | -0.152 | 0.0000 |  | 0.0000 | 0.0000 | * | |  | * | * |
| jul_temp_max | -1.086 | -0.430 | -0.405 | -1.086 | -0.342 | 0.103 | 0.052 | 0.159 | 0.9697 | 0.0000 | 0.9697 | 0.0005 |  | | * |  | * |
| jul_temp_min | 1.849 | 1.123 | 1.501 | 1.849 | 1.003 | 0 | 0 | -0.100 | 0.2139 | 0.6334 | 0.2139 | 0.0127 |  | |  |  | * |
| jun_rain_tot | -0.064 | -0.266 |  | -0.064 | -0.225 | -0.108 | -0.010 | -0.104 | 0.0142 |  | 0.0142 | 0.0145 | * | |  | * | * |
| jun_temp_max | 1.293 | 0.430 | 1.269 | 1.293 | 0.435 | 0 | 0 | 0.068 | 0.0134 | 0.1631 | 0.0134 | 0.2878 | * | |  | * |  |
| jun_temp_min | -2.219 | -0.747 | -1.949 | -2.219 | -0.774 | 0 | 0 | 0.072 | 0.3766 | 0.3315 | 0.3766 | 0.3515 |  | |  |  |  |
| mar_rain_tot | -0.234 |  | -0.176 | -0.234 |  | 0 | 0 | 0 | 0.1118 | 0.0193 | 0.1118 |  |  | | * |  |  |
| mar_temp_max | 0.523 | 0.378 | 0.238 | 0.523 | 0.355 | 0.006 | 0 | 0 | 0.1178 | 0.1307 | 0.1178 | 0.3311 |  | |  |  |  |
| mar_temp_min | 0.466 |  | 0.794 | 0.466 | -0.038 | 0 | 0 | -0.017 | 0.7155 | 0.0333 | 0.7155 | 0.7727 |  | | * |  |  |
| may_rain_tot | 0.426 | 0.134 | 0.279 | 0.426 |  | 0 | 0 | -0.102 | 0.8492 | 0.0179 | 0.8492 |  |  | | * |  |  |
| may_temp_max | 0.173 |  |  | 0.173 | -0.099 | 0 | 0 | -0.079 | 0.4364 |  | 0.4364 | 0.0210 |  | |  |  | * |
| may_temp_min | -0.969 | -0.524 | -0.845 | -0.969 | -0.447 | -0.035 | 0 | -0.087 | 0.1127 | 0.0000 | 0.1127 | 0.0787 |  | | * |  |  |

**Table S2**: **Estimated coefficients and their p-values and significance for 7 variable selection methods**, as well as the full climate model (c.m.). cv.b = best subset selection with cross-validation (Method 1), s.bo = cross stepwise selection in both backwards and forwards directions (Method 2a), s.ba = 10-fold cross-validation backwards stepwise selection (Method 2b), s.f = 10-fold cross-validation forwards stepwise selection (Method 2c), l.m = cross-validation lasso using lambda that minimises the prediction error (Method 3a), l.1 = cross-validation lasso model using lambda for smallest model and within 1 standard error (Method 3b), e.n = cross-validation elastic net using lambda and alpha that minimises the prediction error (Method 4). Of the 7 models shown, only 4 give p-values and significance of each variable.

**Table S3: Adjusted R^2^ and RMSE for the 7 variable selection methods**, as well as the full climate model (c.m.). cv.b = best subset selection with cross-validation (Method 1), s.bo = cross stepwise selection in both backwards and forwards directions (Method 2a), s.ba = 10-fold cross-validation backwards stepwise selection (Method 2b), s.f = 10-fold cross-validation forwards stepwise selection (Method 2c), l.m = cross-validation lasso using lambda that minimises the prediction error (Method 3a), l.1 = cross-validation lasso model using lambda for smallest model and within 1 standard error (Method 3b), e.n. = cross-validation elastic net using lambda and alpha that minimises the prediction error (Method 4).

| **Model** | **Adjusted R^2^** | **RMSE** |
| --- | --- | --- |
| c.m | 0.449 | 0.376 |
| **cv.b** | **0.466** | **0.383** |
| **s.bo** | **0.459** | **0.384** |
| s.ba | 0.449 | 0.451 |
| s.f | 0.443 | 0.446 |
| l.m | -0.065 | 0.477 |
| l.1 | 0.024 | 0.496 |
| e.n | -1.08 | 0.428 |

***Table S4: The 6 main principal components contributing to variance in spring barley yields***. The proportion of variance of each PC is shown, along with the degree of correlation of each climate variable with each PC. Variables correlating by more than |0.3| are in bold.

|  | | **Principal Component** | | | | | |
| --- | --- | --- | --- | --- | --- | --- | --- |
|  |  | 1 | 2 | 3 | 4 | 5 | 6 |
| **Proportion of variance** | | **0.26** | **0.22** | **0.18** | **0.10** | **0.091** | **0.043** |
| Climate variable | Apr_rain_tot | 0.24 | 0 | -0.03 | **0.44** | -0.25 | 0.08 |
|  | Aug_rain_tot | 0.19 | **0.34** | 0.1 | -0.09 | 0.26 | 0.16 |
|  | Jul_rain_tot | -0.11 | -0.16 | 0.3 | **-0.4** | -0.04 | -0.09 |
|  | Jun_rain_tot | 0.1 | -0.15 | **0.32** | -0.03 | -0.24 | **0.53** |
|  | Mar_rain_tot | 0.22 | 0.23 | 0.24 | -0.29 | 0.07 | **0.33** |
|  | May_rain_tot | **-0.35** | -0.18 | -0.09 | -0.17 | 0.12 | 0.25 |
|  | Apr_rain_dmax | 0.03 | -0.24 | 0.19 | **0.47** | -0.24 | 0.23 |
|  | Apr_temp_max | -0.2 | 0 | -0.26 | -0.11 | **-0.46** | -0.21 |
|  | Aug_temp_max | **-0.38** | -0.02 | -0.2 | 0.16 | 0.06 | 0.3 |
|  | Jul_temp_max | 0.03 | **0.43** | -0.14 | 0.2 | -0.01 | 0.12 |
|  | Jun_temp_max | -0.2 | 0.29 | -0.14 | 0.09 | **0.35** | 0.05 |
|  | Mar_temp_max | 0.17 | -0.02 | 0.28 | **0.31** | **0.38** | -0.26 |
|  | May_temp_max | 0.12 | **0.38** | -0.11 | -0.03 | -0.24 | 0.12 |
|  | Apr_temp_min | **0.35** | -0.06 | -0.17 | -0.22 | -0.21 | -0.23 |
|  | Aug_temp_min | -0.03 | -0.27 | **-0.4** | 0.05 | 0.19 | 0.24 |
|  | Jul_temp_min | **0.35** | 0.07 | **-0.31** | -0.02 | -0.08 | 0.11 |
|  | Jun_temp_min | 0.24 | -0.17 | **-0.35** | 0.03 | 0.22 | -0.03 |
|  | Mar_temp_min | 0.24 | **-0.36** | 0.05 | 0.07 | 0.21 | -0.07 |
|  | May_temp_min | 0.28 | -0.21 | -0.21 | -0.25 | 0.12 | 0.3 |

**Table S5**: **Statistical significance and corresponding coefficient of each climate variable in the mixed model** with year, variety and year:farm. Significant variables are shown in **bold**. The AIC of the overall model is given, with lower values corresponding to a better model fit.

| **Climate variable** | **Significance in model** | **Coefficient** | **AIC** |
| --- | --- | --- | --- |
| - | - | - | 127.5 |
| Jul_temp_max | **0.004** | 0.27 | 123.6 |
| Aug_temp_max | **0.024** | 0.20 | 127.4 |
| Jul_rain_tot | **0.028** | -0.0069 | 135.0 |
| Apr_temp_min | 0.059 | -0.097 | 130.0 |
| Jun_temp_min | 0.090 | -0.11 | 130.4 |
| May_temp_min | 0.101 | -0.10 | 130.5 |
| Mar_temp_min | 0.110 | -0.095 | 130.7 |
| Jun_temp_max | 0.120 | 0.11 | 130.6 |
| Jun_rain_tot | 0.131 | -0.0037 | 137.4 |
| Mar_rain_tot | 0.190 | -0.0035 | 137.7 |
| May_temp_max | 0.212 | 0.098 | 131.2 |
| Jul_temp_min | 0.340 | -0.075 | 131.9 |
| May_rain_tot | 0.534 | 0.0034 | 137.8 |
| Aug_temp_min | 0.540 | -0.044 | 132.5 |
| Apr_rain_tot | 0.573 | 0.058 | 132.0 |
| Mar_temp_max | 0.591 | -0.0020 | 138.6 |
| Aug_rain_tot | 0.635 | 0.062 | 131.5 |
| Apr_rain_dmax | 0.693 | -0.0013 | 139.0 |
| Apr_temp_max | 0.842 | -0.026 | 131.7 |

References

Friedman, J., Hastie, T., & Tibshirani, R. (2010). Regularization Paths for Generalized Linear Models via Coordinate Descent. *Journal of Statistical Software*, **33**(1), 1–22.

Kuhn, M. (2020). caret: Classification and Regression Training. Retrieved from https://cran.r-project.org/package=caret

Lumley, T. (2020). leaps: Regression Subset Selection. Retrieved from https://cran.r-project.org/package=leaps

Venables, W. N., & Ripley, B. D. (2002). *Modern Applied Statistics with S*, Fourth, New York: Springer.

Wickham, H., Averick, M., Bryan, J., … Yutani, H. (2019). Welcome to the Tidyverse. *Journal of Open Source Software*, **4**(43). doi:10.21105/joss.01686
